# A temporal allocation of amino acid resources ensures fitness and body allometry

**DOI:** 10.1101/2023.12.08.570599

**Authors:** Luca Valzania, Aya Alami, Pierre Léopold

## Abstract

Organisms have evolved strategies to store resources and overcome periods of low or no nutrient access including transient shortages or longer non-feeding developmental transitions. Holometabolous insects like *Drosophila* represent an attractive model to study resource allocation during development since they alternate feeding and non-feeding periods. Amino acids are essential components for tissue growth and renewal, but the strategies used for their storage remain largely unexplored. Here, we uncover a 2-step mechanism for the temporal production, storage and use of specific storage proteins called hexamerins. Moreover, we show that modulating amino acid storage affects the relative growth of organs across developmental phases and, as a consequence, alters body allometry and adult fitness.

## INTRODUCTION

All organisms rely on nutrient stores during the course of their lives. For the vast majority of animal species, embryos do not have the ability to feed during early stages of their development and rely on maternal reserves stored in the egg ^1^. Animals that hibernate or those that make long migrations do not feed for prolonged periods and use nutrient reserves they have previously accumulated during periods of food availability ^2,3^. Storage availability is also a determinant of the social identity, as seen in the case of vitellogenin for cast determination in social insects ^4^. Finally, occasional food shortages or competition for food within a population also force individuals to use stored resources. Therefore, an effective orchestration of resource allocation across the life-course of an individual is crucial for evolutionary success, yet the mechanisms remain largely unknown.

Development of holometabolous insects provides an interesting paradigm in which to approach these questions ^5^. Indeed, the life cycle of these insects alternates non-feeding or “closed” period (embryonic and pupal) with feeding or “open” ones (larval and adult). The resources required to proceed through a closed developmental phase therefore come from the stores made during the preceding open phase.

So far, studies on nutrient and energy storage across animal species have largely focused on fat and carbohydrates ^6–8^, with protein and amino-acid stores remaining largely under-investigated.

Dedicated proteins that serve as reserves of metal ions and amino acids are present in plants and animals and function as important resources for developing organisms ^9,10^. The best characterized storage proteins are the insect larval serum proteins (LSPs) or hexamerins, a family of proteins that circulate in the hemolymph as hexamers of different subunits ^11,12^. They are synthesized in large amounts by the fat body (functional homolog of the vertebrate liver and fat) of feeding larvae and secreted into the hemolymph, where they represent up to 90% of the total circulating proteins ^13^. However, despite their potential role in resource allocation, the mechanistic and developmental aspects of amino-acid storage played by hexamerins remain poorly characterized.

Here, we use *Drosophila* as a powerful molecular genetic model for studying the dynamic allocation of proteins and their constituent amino acids. We identify the major players in the process of protein storage and their fluxes operating across developmental periods. We bring an *in vivo* demonstration that the formation of a circulating pool of hexamerins in the larval hemolymph is under the control of nutrition through TOR signalling. We further show that the temporally controlled hexamerin flux from the hemolymph back to the fat body relies on a specific serum protein called FBP1 and of an ecdysone timer operating in fat body cells. Finally, we bring evidence that this temporal amino-acid allocation is essential to ensure adult fitness and organ-to-body allometry.

## RESULTS

### Storage protein expression is restricted to the fat body and relies on nutritional and developmental inputs

In *Drosophila*, storage proteins called hexamerins consist of 4 Larval Serum Proteins (LSP1α, β, γ, LSP2) showing a high sequence similarity ^11^. In addition, Fat Body Protein 1 and 2 (FBP1, FBP2) have been proposed as putative receptors for LSP endocytosis into the fat body ^14^. We first sought to determine the precise timing and tissue source for the production of these components. q-RT PCR analysis revealed a peak of expression at the end of the larval feeding period for each *Lsp* and *Fbp2* gene, while *Fbp1* showed a slightly delayed expression peaking during the very early prepupal phase (Fig. 1A-D). Silencing each of these genes specifically in the fat body using the *Lpp-Gal4* driver fully suppressed their expression (Fig. S1A-F), indicating that the fat body is the only source for their production. In addition, silencing any of the *Lsp1* genes (α, β or γ) drastically decreased the expression of other *Lsp1* genes, indicative of a co-regulation of all LSP1 subunits (Fig. S1G-I). In contrast, downregulating the expression of *Lsp2* induced the upregulation of all three *Lsp1* genes (Fig. S1J), revealing a compensatory mechanism maintaining global circulating LSP levels. These co-regulations were not observed upon *Fbp1* or *Fbp2* silencing (Fig. S1L, M), even if the expression of the two putative receptors decreased in absence of all *Lsp* (Fig. S1K).

**Figure 1.**
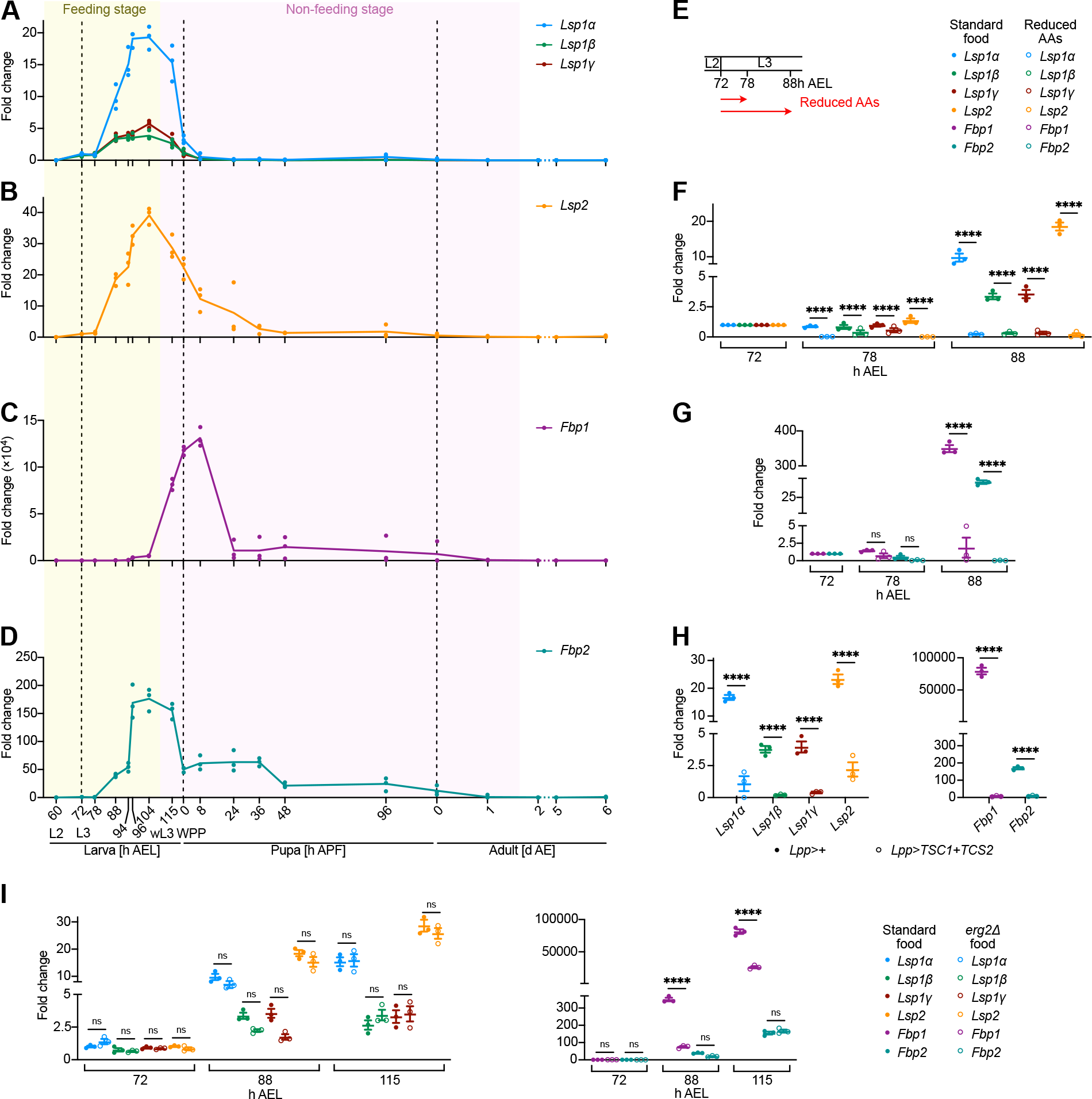
Developmental and environmental control of *Lsp* and *Fbp* expression. (A-D) mRNA levels of *Lsp1*α, *Lsp1*β, *Lsp1*γ (A), *Lsp2* (B), *Fbp1* (C), and *Fbp2* (D) during the development, measured by RT-PCR. (E) Schematic representation of amino acid restriction experiments. Larvae were shifted to low amino acid food at 72h AEL. (F-G) mRNA levels of *Lsp* and *Fbp* genes in larvae reared on standard food or in presence of a reduced amount of amino acids (Mann-Whitney unpaired nonparametric t test. ns: not significant; ****p≤0.0001). (H) RT-PCR of *Lsp* and *Fbp* genes from larvae in which TOR pathway was specifically downregulated in the fat body (*Lpp>TSC1+TSC2*) and from control larvae (*Lpp>+*). Larvae were reared at 18ºC until 52h AEL and then transferred to 25ºC. Samples were collected when larvae were in their late wandering stage (Mann-Whitney unpaired nonparametric t test. ****p≤0.0001). (I) mRNA levels of *Lsp* and *Fbp* genes measured by RT-PCR in larvae reared on standard food or on *erg2*Δ food (Mann-Whitney unpaired nonparametric t test. ns: not significant; ****p≤0.0001). In all graphs represented in Figure 1, results shown are mean±SEM and n=3 replicates per time point. Fold changes are normalized by *rp49*, and developmental time points are expressed in hours after egg laying [h AEL], hours after pupa formation [h APF] and days after eclosion [d AE].

We next examined the basis for the temporal regulation of *Lsp* and *Fbp* gene expression. Hexamerins are produced during the feeding period. Reducing amino-acid content in the food starting early in the 3^rd^ instar drastically reduced expression of all *Lsp* and *Fbp* genes (Fig. 1E-G). The Target Of Rapamycin (TOR) signaling pathway plays a major role in sensing nutritional inputs through the presence of amino acids in fat body cells ^15,16^. Mimicking an amino-acid restriction by suppressing TORC1 activity specifically in fat cells (*Lpp>TSC1+TSC2*) abrogated expression of all *Lsp* and *Fbp* genes (Fig. 1H). Since all storage proteins – encoding genes are transcribed at the transition between larva and pupa, we decided to investigate the role of ecdysone, whose levels peak during this period.

Placing animals on *erg2Δ* mutant yeast food (Fig. S1N and Methods, ^17^) prevented the peak of ecdysone accumulation at the larva-to-pupa (L/P) transition and abrogated expression of *Fbp1*, but none of the *Lsp* genes nor *Fbp2* were affected (Fig. 1I). Taken together, our genetic analysis established that hexamerins and their putative receptors are produced by fat cells before the L/P transition and that nutrition is a major driver for their expression, mediated by adipose TOR signaling. Accordingly, *Lsp* expression declines early during early pupa after animals stop feeding. Importantly, our data reveal that, in addition to the requirement for nutrition, *Fbp1* expression is timed by a developmental signal. Indeed, the rise of ecdysone marking the end of the larval phase activates its transcription, suggesting a specific role for the FBP1 protein in the temporal orchestration of amino acid resources.

### LSPs and FBPs are secreted into the hemolymph and their uptake is mediated by FBP1

While produced in fat cells, hexamerins are proposed to be released in the open circulation, forming an important store of proteins circulating at high concentrations in the hemolymph ^13^. To understand the dynamics of accumulation of circulating hexamerins, we generated specific antibodies against LSP1α, LSP2, FBP1 and FBP2 (see Methods; Fig. S2A-D). As expected, LSP1α and LSP2 were present in the hemolymph of late larvae. Surprisingly, both FBP1 and FBP2 were also found circulating in the hemolymph in contrast to their previously suggested role as membrane-associated receptors (Fig. 2A, B). This, together with the absence of trans-membrane domain signature in these proteins, called for a reinterpretation of their function and suggested that both FBPs may serve as circulating storage proteins *in vivo*. Concerning FBP1, western blotting analysis revealed that only a processed 64 kDa form was present in the hemolymph, while larger and shorter forms were also found in fat body cells (Fig. 2A). To identify the circulating hexamerin complexes in more details, we performed native gel analysis on hemolymph extracts followed by western blotting analysis against the various LSPs and FBPs. The co-labelling of discrete bands with the different antibodies suggests that LSPs and FBPs could associate in various complexes *in vivo*, all potentially containing FBP1 (Fig. 2C). This also suggests that FBP1 could be secreted after proteolytic cleavage to form circulating complexes with other LSPs. To better elucidate the relationship between FBP1 and LSPs, we analyzed the various amounts of LSP1α, LSP2 and FBP2 in the hemolymph and in the fat body both in control conditions and after blocking expression of *Fbp1* in the fat body by RNAi-mediated silencing. In control conditions, LSPs and FBP2 disappear from the hemolymph and concomitantly accumulate in fat cells at the L/P transition, suggestive of a re-uptake by FB cells. In contrast, silencing *Fbp1* maintained high levels of LSPs and FBP2 in the hemolymph and reduced their amount in the fat body after the L/P transition (Fig. 2D-I, Fig. S2E). This suggested that FBP1 may have a crucial role in the reuptake of LSPs and FBP2 in fat cells at the L/P transition. To confirm the relevance of FBP1 in this process, we used the *QF2/QUAS* system to ectopically express HA-tagged forms of LPS1α or LSP2 in larval muscles, a tissue that does not produce hexamerins in physiological conditions. This experimental setup allows visualizing the uptake of HA-tagged LSP1α and LSP2 in fat cells, separately from their endogenous production by fat body cells (see rationale in Fig. 2J). We observed that HA-tagged LSP1α and LSP2 produced in muscle cells are effectively secreted in the hemolymph where they form complexes comparable in size to the complexes formed by endogenous LSP1α and LSP2 (Fig. 2K, L). Strikingly, suppression of *Fbp1* expression in fat body cells (*Lpp>Fbp1*^*RNAi*^) prevented circulating LSP1α-HA and LSP2-HA to enter fat body cells (Fig. 2M). This was not observed when *Lsp1α, Lsp2* or *Fbp2* were silenced (Fig. 2M). In addition, no LSP1α-HA or LSP2-HA uptake was observed in fat cells before *Fbp1* expression starts at the time of L/P transition (Fig. S2F, 94h AEL). This and our previously described results demonstrate that circulating LSP1α and LSP2 complexes re-enter the fat body at the L/P transition and that FBP1 is specifically required for this process.

**Figure 2.**
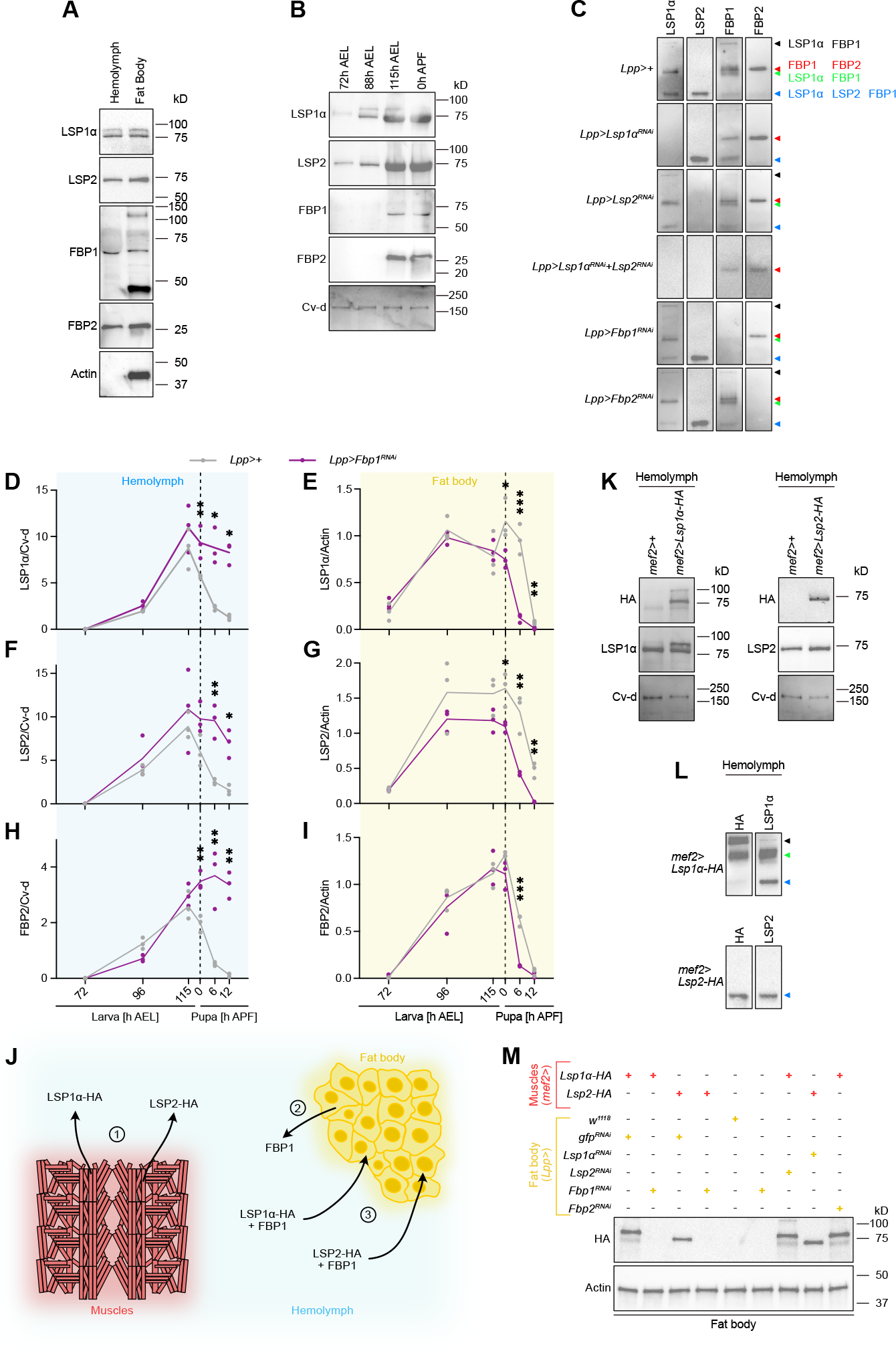
LSPs and FBPs are secreted into the hemolymph and re-enter the fat body through FBP1. (A) Immunoblots of hemolymph and fat body extracts from late wandering larvae. Samples were probed with antibodies to LSP1α, LSP2, FBP1, FBP2, and Actin, with molecular masses indicated to the right. (B) Immunoblots of hemolymph extracts from L3 larvae and pupae. Samples were collected at different time points indicated above the panel, and were probed with antibodies against LSP1α, LSP2, FBP1, FBP2, and Cv-d as a loading control. References for molecular masses are indicated to the right of each blot. (C) Native western blots of hemolymph extracts from *Lpp>+, Lpp>Lsp1*α^*RNAi*^, *Lpp>Lsp2*^*RNAi*^, *Lpp>Lsp1*α^*RNAi*^*+Lsp2*^*RNAi*^, *Lpp>Fbp1*^*RNAi*^, and *Lpp>Fbp2*^*RNAi*^ larvae. Samples were collected at 115h after egg laying and probed with antibodies to LSP1α, LSP2, FBP1, and FBP2. Colored arrowheads and the present proteins distinguish the different complexes on the right of each blot. (D-I) Quantification of LSP1α, LSP2, and FBP2 levels in hemolymph or fat body cells of *Lpp>+* or *Lpp>Fbp1*^*RNAi*^ animals. Protein levels are normalized by Cv-d in hemolymph and Actin in fat body. Results shown are mean±SEM and n=3 replicates per time point (Mann-Whitney unpaired nonparametric t test. *p≤0.05, **p≤0.01, ***p≤0.001). Original western blots are shown in Supplemental Figure 2E. (J) Schematic representation of LSPs-HA experimental approach. *Lsp1α-HA* and *Lsp2-HA* are expressed in larval muscles and secreted into hemolymph (1); endogenous FBP1 is expressed in fat body, secreted into hemolymph and able to bind LSP1α-HA and LSP2-HA (2). LSPs-FBP1 complexes can be taken-up by the fat body (3). (K) Immunoblots of hemolymph extracts from *mef2>+, mef2>Lsp1α-HA*, and *mef2>Lsp2-HA* larvae. Samples were collected at 115h AEL and probed with antibodies to HA, LSP1α, LSP2, and Cv-d. References for molecular masses are indicated to the right of each blot. (L) Native western blots of hemolymph extracts from *mef2>Lsp1α-HA*, and *mef2>Lsp2-HA* larvae. Samples were collected at 115h AEL and probed with antibodies to HA, LSP1α, and LSP2. Colored arrowheads, on the right of each blot, distinguish the different complexes with the same color code used in Figure 2C. (M) Immunoblots of fat body extracts from animals expressing different transgenes in muscles (indicated in red) and in fat body cells (reported in yellow). Samples were probed with antibodies against HA and Actin, with molecular masses indicated to the right.

Amino acid storage thus appears to be regulated at two levels. At first, the presence of nutrients regulates hexamerin expression, then entry in pupal development, which marks the start of the closed period, determines when LSPs can be taken up by the fat body through the production of FBP1.

### LSP production is an energy-costing process during larval development

We then genetically impaired each of the two levels of regulation to test the role of protein storage during development. For this, we measured different larval parameters (Fig. S3A) in control, no LSP production (*Lpp>Lsp1α*^*RNAi*^*+Lsp2*^*RNAi*^, hereafter referred to as “pan-*Lsp* silencing”) or no LSP uptake (*Lpp>Fbp1*^*RNAi*^) conditions. Intriguingly, larval wet mass was significantly increased both upon pan-*Lsp* or *Fbp1* silencing (Fig. 3A), similarly to pupal volume (Fig. S3D). However, dry mass was only increased in pan-*Lsp* silencing (Fig. 3C), whereas *Fbp1* silencing led to an increase in hemolymph volume (Fig. 3B, Fig. S3B). Therefore, preventing the formation of LSP stores led to a general increase in larval tissue mass at pupariation, whereas preventing LSPs from re-entering the fat body led to a modification of larval water balance, which could be explained by an increase in osmotic pressure due to a large quantity of LSP proteins accumulating in the serum. Remarkably, pan-*Lsp*-silenced larvae pupariated earlier that control and *Fbp1*-silenced ones (Fig. 3D, Fig. S3C), suggesting that all features of development were accelerated in these conditions. Accordingly, the volume of imaginal wing discs at the white prepupal stage was significantly increased (Fig. 3E, F) indicative of an acceleration in tissue growth in pan-*Lsp* silencing conditions. Feeding activity was similar in pan-*Lsp* silencing and control conditions, indicating that acceleration of growth was not due to an increase in nutritional input (Fig. 3G). In line with this, the level of phosphorylation of S6 kinase, a marker of TORC1 activity, in fat tissue was identical in the three conditions (Fig. 3H, I). Altogether, our data suggest that constituting a large stock of amino acids in the form of hexamerins at the end of larval development is an energy-costing process that has a trade-off for the growth of early developing adult structures like the wing imaginal discs.

**Figure 3.**
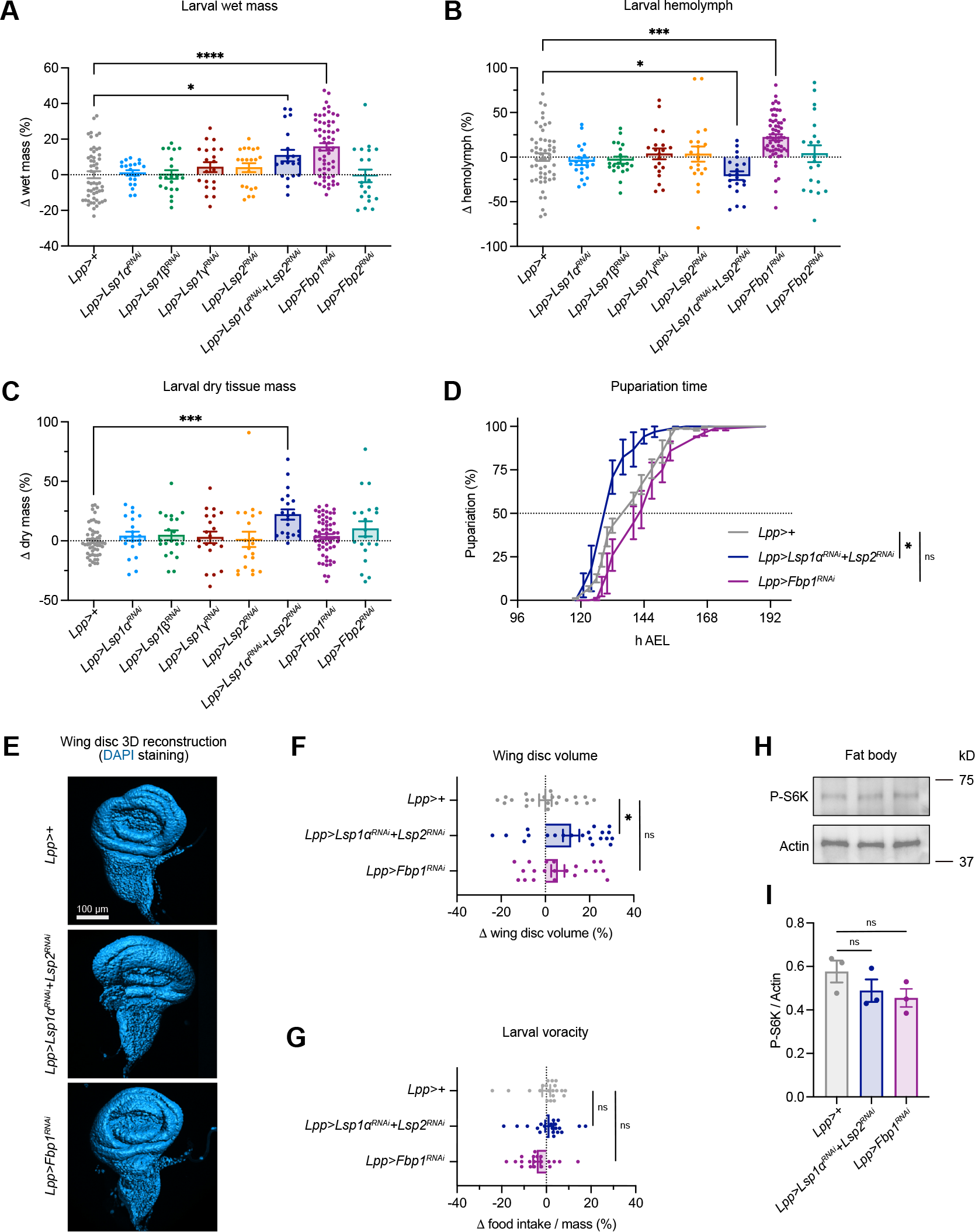
Suppression of hexamerin storage enhances larval growth. (A) Quantification of larval wet mass with the indicated genotypes (n=20-58 for each genotype). Results shown are mean±SEM as percentage difference compared to control *Lpp>+* (one-way ANOVA followed by Dunnem’s multiple-comparisons test. Larvae of all genotypes were compared to control, but only the significantly different pairs are reported in the graph. *p≤0.05, ****p≤0.0001). (B) Differences in larval hemolymph volume with the indicated genotypes (n=20-58 for each genotype). In each column mean±SEM of the delta between *Lpp>+* and the corresponding genotype is plotted (one-way ANOVA followed by Dunnem’s multiple-comparisons test. Only significant different pairs are shown in the graph. *p≤0.05, ***p≤0.001). (C) Variations in larval dry tissue mass are represented in the graph (n=20-58 for each genotype). Percentage values are the difference between *Lpp>+* control larvae and the analyzed genotypes (one-way ANOVA followed by Dunnem’s multiple-comparisons test. Pairs showing significant difference are indicated in the graph. ***p≤0.001). (D) Pupariation timing of *Lpp>+, Lpp>Lsp1α*^*RNAi*^*+Lsp2*^*RNAi*^, and *Lpp>Fbp1*^*RNAi*^ (one-way ANOVA followed by Dunnem’s multiple-comparisons test. ns: not significant; *p≤0.05). (E) Representative examples of surface reconstruction for volume measurements of the wing disc stained with DAPI at white prepupal stage. Scale bar is the same for the three images and corresponds to 100μm. (F) Quantification of the wing disc volume difference between the control *Lpp>+, Lpp>Lsp1α*^*RNAi*^*+Lsp2*^*RNAi*^ and *Lpp>Fbp1*^*RNAi*^ discs (n=21 per genotype. One-way ANOVA followed by Dunnem’s multiple-comparisons test. ns: not significant; *p≤0.05). (G) Food intake is measured using the blue dye Erioglaucine in the food and normalised to the body weight (n=20 larvae for each genotype). Percentage values are the difference between *Lpp>+* control larvae and the analyzed genotypes (One-way ANOVA followed by Dunnem’s multiple-comparisons test. ns: not significant). (H) Immunoblot of fat body extracts from *Lpp>+, Lpp>Lsp1α*^*RNAi*^*+Lsp2*^*RNAi*^ and *Lpp>Fbp1*^*RNAi*^ larvae. Samples were probed with antibodies against P-S6K and Actin, with molecular masses indicated to the right. (I) Quantification of the bands from the above western blot analysis (One-way ANOVA followed by Dunnem’s multiple-comparisons test. ns: not significant).

### A deficit in LSP production or reuptake affects adult fitness and body proportions

After the larval phase, animals undergo metamorphosis, a closed phase of development involving important organogenesis in absence of nutrient intake. We therefore investigated the role of LSPs and FBPs in providing amino-acid resources needed during this phase. Consistent with a function of hexamerins in nutrient storage, preventing LSP production (pan-*Lsp* silencing) or their reuptake in the fat body (*Fbp1* silencing) had a strong effect on pupal and adult development. Both conditions generated a partial lethality during pupa and more than 30hrs delay at eclosion for surviving animals (Fig. 4A). Emerging adults were analyzed for locomotion, fertility, starvation survival, and longevity.

**Figure 4.**
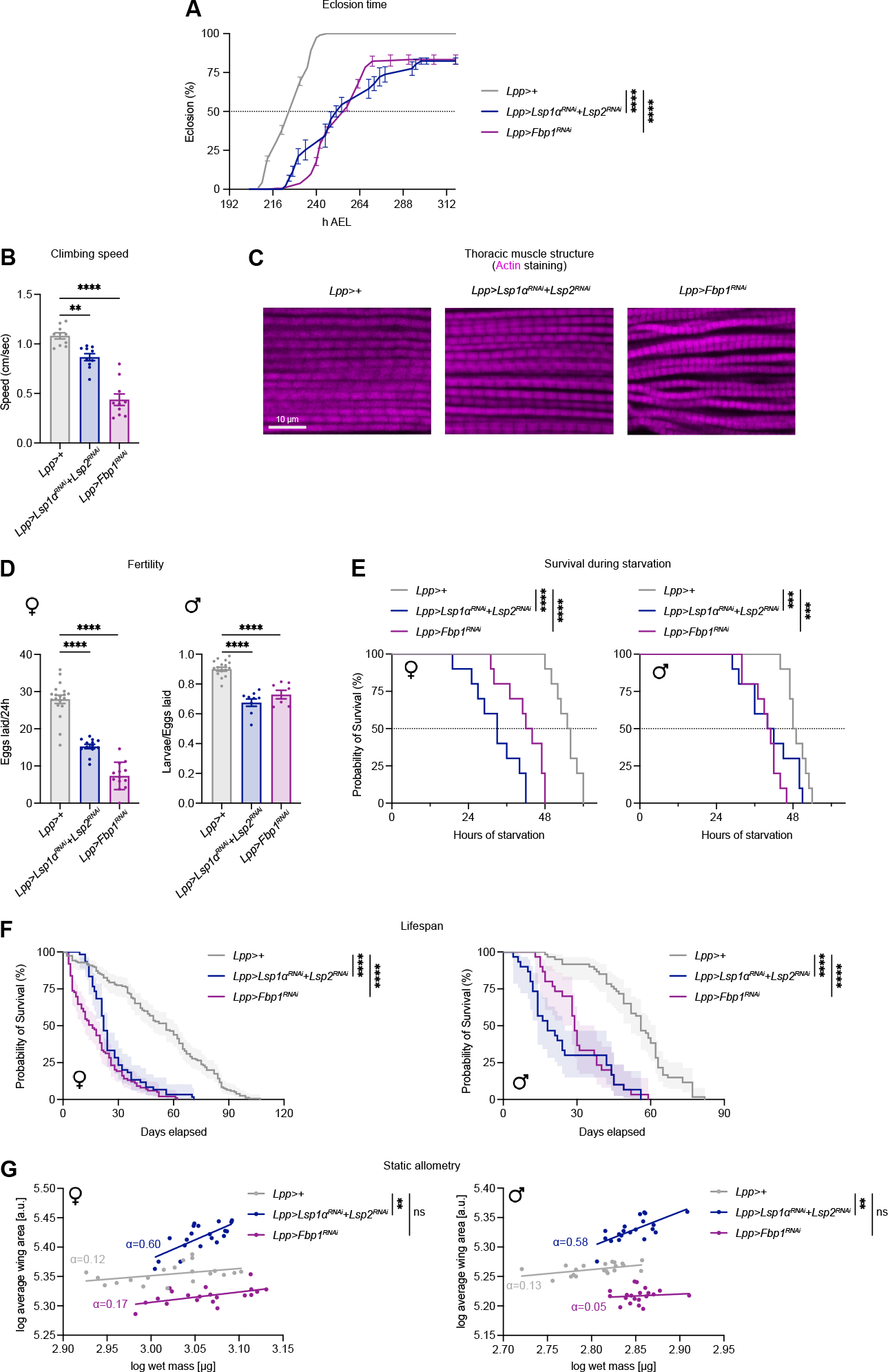
Absence of hexamerins during pupal development deteriorates adult fitness. (A) Eclosion timing of *Lpp>+, Lpp>Lsp1α*^*RNAi*^*+Lsp2*^*RNAi*^ and *Lpp>Fbp1*^*RNAi*^ adults. Results are shown as means with SEM at different time points, connected by lines; n=3 represents average of three vials per genotype in one experiment (one-way ANOVA followed by Dunnem’s multiple-comparisons test. ****p≤0.0001). (B) Climbing speed of *Lpp>+, Lpp>Lsp1α*^*RNAi*^*+Lsp2*^*RNAi*^ and *Lpp>Fbp1*^*RNAi*^ flies (n=10 means the average of ten different measurements of climbing speed for ten groups of flies for each genotype). Results are mean±SEM (one-way ANOVA followed by Dunnem’s multiple-comparisons test. **p≤0.01, ****p≤0.0001). (C) Confocal images of thoracic muscles, stained with Phalloidin, from *Lpp>+, Lpp>Lsp1α*^*RNAi*^*+Lsp2*^*RNAi*^ and *Lpp>Fbp1*^*RNAi*^ flies. Scale bar is equal in all images and corresponds to 10μm. (D) Tests on *Lpp>+, Lpp>Lsp1α*^*RNAi*^*+Lsp2*^*RNAi*^ and *Lpp>Fbp1*^*RNAi*^ flies to assess the fertility. In the lej graph female fertility was evaluated by counting the number of eggs laid by a female in 24h. On the right graph male fertility was estimated by the ratio between larvae and eggs laid. Results in both graphs are shown as mean±SEM; n=10-19 means the average of eggs laid or hatched in 10-19 groups of flies for each genotype (one-way ANOVA followed by Dunnem’s multiple-comparisons test. ****p≤0.0001). (E) Survival curves of adult females (lej graph) and males (right graph) being reared on starved conditions after their eclosion. Results are shown in percent staircase graph; n=10 animals (one-way ANOVA followed by Dunnem’s multiple-comparisons test. ***p≤0.001, ****p≤0.0001). (F) Survival curves of adult females (lej graph) and males (right graph) reared on standard food with the corresponding genotypes (n=60-155 females per genotype and 30-60 males for each genotype). Results are present as percent staircases with 95% confidence interval (Log-rank (Mantel-Cox) and Gehan-Breslow-Wilcoxon tests. ****p≤0.0001). (G) The scaling relationship for wing size against body size was obtained by transforming raw data in logarithm and plotted in log-log graph; n=20 females and 20 males per genotype (equations for simple linear regression and tests for slope and intercept significancy were generated by GraphPad Prism. ns: not significant; **p≤0.01). Allometric coefficients (α) of different lines are reported in the graphs.

Both genetic conditions affected all traits of adult fitness, including reduced climbing ability associated with defects in muscle structure (Fig. 4B, C), reduced survival to starvation post-emergence (Fig. 4E) and severely reduced lifespan (Fig. 4F). Fertility of males was also reduced and more severely for females (Fig. 4D). Finally, adult mass was slightly increased in both conditions (Fig. S4A). Again, this increase was due to dry mass increase in the case of pan-*Lsp* knock-down and hemolymph volume increase in the case of *Fbp1*^*RNAi*^ (Fig. S4B, C). Notably, pan-*Lsp* knock-down adult only showed a 6% increase in dry mass (Fig. S4C), to compare with a 22% increased larval dry mass, demonstrating the need in larval protein stores during pupal stage for proper development of adult structures. Accordingly, adult dry mass was reduced by 10% upon *Fbp1* silencing (Fig. S4C). Adult wing area was increased in pan-*Lsp* knock-down (Fig. S4D), as a consequence of increased larval disc growth (see Fig. 3E, F). In contrast, *Fbp1*^*RNAi*^ flies showed reduced wing area (Fig. S4D), indicative of a reduced expansion of imaginal tissues during the pupal period due to limited resources. These results reveal two combined effects of preventing the formation of hexamerin stores during late larval life: while larval tissues and early forming imaginal structures are favored, late pupal structures are affected. By contrast, blocking LSP reuptake by fat cells precludes their use during pupa and only affects late forming imaginal structures.

These contrasting developmental defects could have differential consequences on the development of adult structures and thus perturb body-organ allometry. We tested this possibility by measuring the ratio between adult wing surface and body weight in different conditions of LSP perturbation. Preventing LSP reuptake in fat cells (*Lpp>Fbp1*^*RNAi*^) had no effect on the allometric relationship between adult wings and body. By contrast, pan-*Lsp* silencing during late larval life (*Lpp>Lsp1α*^*RNAi*^*+Lsp2*^*RNAi*^) strongly modified the allometric parameters between body weight and wing surface (Fig. 4G, Fig. S4E). Overall, these results demonstrates that protein storage is made at a cost for the early development of imaginal tissues during larval life, while contributing to the formation of future adult structures during pupa. This illustrates how protein storage contributes to the precise temporal allocation of amino acid resources during development, ensuring allometric growth and fitness of emerging adults.

## DISCUSSION

In this work, we investigate through genetic and biochemical analysis the role of amino acid resources and their proper temporal allocation during *Drosophila* development. Contrarily to previous conclusions, our work establishes that LSP1, LSP2 but also FBP1 and FBP2 contribute to the circulating pool of hexamerins and that their production by fat body cells relies on TOR pathway in response to favorable nutritional input. TOR signaling in fat cells also controls the level of circulating insulin-like peptides, promoting imaginal tissue growth ^16,18^. Therefore, amino-acid storage and imaginal disc growth are two antagonistic processes that both rely on TOR signaling in fat body cells.

Interestingly, the level of feeding and fat body TOR signaling does not vary in LSP-less conditions, suggesting that in these conditions, TOR function is somehow re-routed towards systemic growth activation. Further work should help better understand the molecular mechanisms ensuring the intracellular balance of TOR signaling between these interdependent functions in fat cells.

Contrarily to previously proposed models, we observed that both FBP1 and FBP2 behave as circulating storage proteins. In addition, we bring functional evidence that FBP1 is required for the reuptake of all hexamerin complexes by the fat body, and that this critical mechanism is timed through the control of *Fbp1* expression by ecdysone at the L/P transition. FBP1 was proposed to be regulated by proteolytic cleavage ^14^. The 64 kDa FBP1 polypeptide we detected in the hemolymph could be the result of ecdysone-dependent proteolytic cleavage. Yet, the FBP1 we noticed accumulating in the fat body after reuptake, measuring 47 kDa, is the result of a second proteolytic cleavage, this time occurred on the secreted FBP1 ^14^. Because FBP1 shares a strong sequence similarity with LSPs ^19^, it is possible that its short isoform accumulating in fat cells may contribute to the pool of stored amino acids like other LSPs.

Finally, since FBP1 is specifically needed for the reuptake of the LSPs by the fat body and it has not domains to link or be part of the cell membrane, the endocytosis of FBP1-containing complexes could necessitate a yet unidentified receptor present at the membrane of fat cells.

The circulating hexamerin pool accounts for more than 10% of the total mass of proteins in late larvae ^20^. Hence, the rapid biosynthesis of this protein pool by the fat body at the end of larval development represents an important metabolic investment by the larva for future pupal development. Indeed, in absence of hexamerin production, we observe that the biosynthetic capacity of the larva is re-routed towards promoting growth of imaginal tissues. This contrasts with our observation that preventing hexamerin reuptake in fat cells during the pupa affects the formation of adult structures and animal fitness. Moreover, it demonstrates the need for storage protein resources during pupal non-feeding development to complete organogenesis and ensure major aspects of adult fitness like reproduction and lifespan. This programmed temporal allocation of amino acids during development is particularly exemplified in our *Lsp*-silencing experiment and its effect on body allometry. Indeed, preventing larval amino acid storage affects early- and late-forming imaginal tissues in opposite ways. Therefore, the proportions of adult body structures are modified. This demonstrates that amino acid resources not only need being properly allocated to peripheral tissues through processing by fat body cells, but that this process needs to be precisely controlled in time to ensure proper organ allometry.

Our study is the first attempt to define the molecular and genetic aspects of protein storage during animal development. It sheds new light on a fascinating process shared by most species to ensure a harmonious growth and fitness despite a highly variable access to amino acid resources during the sensitive period of development.

## MATERIALS AND METHODS

### Drosophila strains and maintenance

All flies were raised, and experiments were conducted using fly food composed of 7.5g/liter agar, 35g/liter wheat flour, 50g/liter yeast powder, 55g/liter sugar, 25ml/liter Methyl, and 4ml/liter propionic acid. The experiments were consistently carried out at a temperature of 25ºC, unless specified in the figure legends. Both male and female flies were utilized in the experiments. Prior to each experiment, a precise staging of the insects was performed by collecting eggs laid over a period of 4h on agar plates with yeast. Newly hatched L1 larvae were gathered after 26h from the start of egg laying and placed in tubes, with each tube accommodating forty larvae and containing standard food. The specific stage of development or the time at which analyses were conducted is indicated in the subsequent sections. *Lpp-Gal4* (Chr X) was a gift from S. Eaton lab ^21^, while *UAS-TSC1+TSC2* was kindly provided by N. Tapon ^22^. The following stocks were obtained from the Vienna Drosophila Research Center (VDRC): *UAS-Lsp1α*^*RNAi*^ (#14898), *UAS-Lsp1β*^*RNAi*^ (#35584), *UAS-Lsp1γ*^*RNAi*^ (#38129), *UAS-Lsp2*^*RNAi*^ (#109979), *UAS-Fbp1*^*RNAi*^ (#37881), and *UAS-Fbp2*^*RNAi*^ (#33172); while the following lines were obtained from the Bloomington Drosophila Stock Center (BDSC) at Indiana University: *w*^*1118*^ (#3605), *Lpp-Gal4* (Chr 3) (#84317), *mef2-QF2* (#66469), and *UAS-gfp*^*RNAi*^ (#44412). *QUAS-Lsp1α-HA* and *QUAS-Lsp2-HA* stocks were generated in this study.

### Quantitative RT-PCR

Larvae were collected at the indicated time point AEL (after egg laying) and frozen immediately in liquid N_2_. Total RNA was extracted using a RNeasy Lipid Tissue Mini Kit (QIAGEN, #74804) according to the manufacturer’s protocol. 1μg of each sample was treated with DNase I (Invitrogen, #18068-015) and reversely transcribed using a SuperScript IV VILO Master Mix (Invitrogen, #11756050). The resulting cDNA was utilized for quantitative qPCR (Viia 7; Applied Biosystems). cDNA was diluted 1:50 in H_2_0 and mixed with 10μM primers and *Power*SYBR Green PCR Master Mix (Applied Biosystems, #4367659). Samples were normalized to ribosomal protein *rp49* transcript levels. Three separate biological samples were collected for each experiment and triplicate measurements were conducted. The following primers were used:

rp49_For: AAGAAGCGCACCAAGCACTTCATC; rp49_Rev: TCTGTTGTCGATACCCTTGGGCTT.

Lsp1α_For: GAGTACATTGCGATGGGAAAGC; Lsp1α_Rev: CATACGAGCGAAGGCCACAT.

Lsp1β_For: GATCGCCATCGCATTGCTG; Lsp1β_Rev: CCCTGCTTGATGTGGTCCT.

Lsp1γ_For: GCCTGTGTGACTGCCTTTAG; Lsp1γ_Rev: AGAGGCTCATCAATACGGTGA.

Lsp2_For: CTTCCAGCACGTCGTCTACTG; Lsp2_Rev: CCCTGCATATCATCACGGAACA.

Fbp1_For: ATCGTGGCGGCATTGATAAGG; Fbp1_Rev: CGAAGGGTGTCAAAGTCCTG.

Fbp2_For: ATGAATCTGACTGGCATGATCCA; Fbp2_Rev: CCAGGCCATAGACAGAGGACA.

### Amino acid restriction assay

Forty precisely staged first instar larvae were grown in a vial containing standard fly flood for 72h AEL then they were transferred on yeast-free medium to reduce the amount of amino acids available in their diet. Larvae were reared on this specific food for 6h or 16h and processed to extract RNA. At least 5 vials were included in the analysis.

### Preparation of *erg2Δ::Trp1* yeast fly food

Fly food made with *erg2Δ::Trp1* yeast was prepared as previously described ^17^. YPH *erg2Δ::Trp1* yeast was grown on a specific medium composed of 6.7g/liter BD Difco™ Yeast Nitrogen Base without Amino Acids (ThermoFisher Scientific, #BD 291940), 5g/liter Bacto™ Casamino Acids (ThermoFisher Scientific, #223050), and 20g/liter Glucose (Sigma-Aldrich, #G7021). 10ml 100X Uracil (Sigma-Aldrich, #U1128) were added and the *erg2Δ::Trp1* yeast was cultured at 29ºC until reaching an OD600=1. The harvested cells were centrifuged and washed with PBS several times, boiled for 30min, and mixed with 0.1g sucrose, 0.12g agar, 10ml H_2_O, and 300μl propionic acid under sterilized conditions. 5ml of this medium were transferred in a vial together with twenty-five sterilized eggs. The tubes were incubated at 25ºC, and larvae were analysed at specific time points.

### Circulating ecdysone measurements

5μl of hemolymph from staged *w*^*1118*^ larvae were homogenized and extracted in 200μl methanol. Samples were processed in a competitive ELISA test as previously published ^23^ to evaluate the amount of circulating 20-E, using the 20-Hydroxyecdysone ELISA kit (Bertin bioreagent, #A05120) as recommended by the manufacturer. Absorbance at 415nm was registered using a TECAN microplate reader.

### Western blots and antibody generation

Three late third instar larvae or fat bodies from three to five animals were homogenized in ice-cold PBS containing Halt™ Protease & Phosphatase Inhibitor Cocktail (ThermoFisher Scientific, #78440) by using a KIMBLE Dounce tissue grinder set (Merck, #D8938), and immediately frozen in liquid N_2_.

Hemolymph samples were prepared by bleeding five animals in a drop of ice-cold PBS containing Halt™ Protease & Phosphatase Inhibitor Cocktail. The collected serum was centrifuged several times at 4ºC to prevent any cells from being present in the final hemolymph samples. Protein content was measured to normalize samples, 4xLaemmli Sample Buffer (Bio-Rad, #1610747) + 10% 2-Mercaptoethanol (Sigma-Aldrich, #M6250) was added, and samples were boiled for 10min at 95°C. Lysates were resolved by electrophoresis using Mini-PROTEAN TGX Precast Gels (Bio-Rad, #4569033) in Tris/Glycine/SDS Buffer (Bio-Rad, #1610732). Proteins were then transferred onto 0.2μm nitrocellulose membranes using the Trans-Blot Turbo Transfer Pack (Bio-Rad, #1704158), blocked in Tris buffered saline (Bio-Rad, #1706435) + 0.1% Tween-20 + 5% BSA, and probed with LSP1α antibody (this study; 1:1000), LSP2 antibody (this study; 1:5000), FBP1 antibody (this study; 1:1000), FBP2 antibody (this study; 1:5000), HA Tag polyclonal antibody (Invitrogen, #71-5500; 1:3000), and Phospho-Drosophila p70 S6 Kinase (Thr398) (P-S6K; Cell Signaling Technology, #9209; 1:1000). For normalization blots were probed with Actin antibody (Sigma-Aldrich, #A2103; 1:5000) or Cv-d antibody (^24^; 1:1000). Horseradish peroxidase (HRP)–conjugated secondary IgG antibodies (Invitrogen, #31470; Jackson ImmunoResearch, #111-035-144; Invitrogen, #A18769; 1:5000) were used together with the Clarity Max™ Western ECL Substrate (Bio-Rad, #1705062) to detect the protein bands. Data show representative results from three biological replicates. Band intensity was quantified using Fiji software. Native western blots were conducted in the same way than standard western blot with only few differences: Native Sample Buffer (Bio-Rad, #1610738) without 2-Mercaptoethanol was added to proteins, samples were immediately run without boiling step, and electrophoresis was performed in Tris/Glycine Buffer (Bio-Rad, #1610734). Polyclonal antibody against LSP1α was generated in rat by the company Eurogentec using the following peptides: YILRENIYEYNQESN (AAs 270-284) and EKYYNYKEYTNYGHF (AAs 787-801). Polyclonal antibody against LSP2 was generated in rat by the company Eurogentec using the following peptides: TFGRNSHYKGSSYVI (AAs 481-495) and VDVKIFHRDEHTNVV (AAs 687-701). Polyclonal antibody against FBP1 was generated in guinea pig by the company Eurogentec using the following peptides: IADSQRYRGGIDKVM (AAs 68-82) and QYQDNNQERDLGQDV (AAs 305-319). Polyclonal antibody against FBP2 was generated in guinea pig by the company Eurogentec using the following peptides: INGEGVLLDKDVETT (AAs 89-103) and FTRSMGDKMIYQKTG (AAs 161-175).

### Generation of transgenic flies

In order to generate the *QUAS-Lsp1*α*-HA* and *QUAS-Lsp2-HA* fly lines, the pQUASamB-Lsp1α-HA and pQUASattB-Lsp2-HA constructs were introduced into the germ line by injections in the presence of the PhiC31 integrase and inserted in the attP40 docking site on the second chromosome (BestGene Inc.). *Lsp1*α*-HA* and *Lsp2-HA* sequences were synthesized by ThermoFisher Scientific. Both sequences have been codon optimized and inserted in pMA-RQ plasmid. These plasmids, in parallel with pQUAST-attB plasmid (Addgene, #104880), were amplified in NEB® Stable Competent *Escherichia coli* (High Efficiency) cells (New England BioLabs, #C3040), purified using NucleoSpin Plasmid kit (MACHENREY-NAGEL, #740588.50) as described in the kit manual, and digested with EcoRI and XbaI (Fastdigest, ThermoFisher Scientific) as reported in their user guide. Lsp1α-HA and Lsp2-HA inserts and pQUAST-attB plasmid were ligated to generate the final pQUASattB-Lsp1α-HA and pQUASattB-Lsp2-HA plasmids.

### Wet mass, hemolymph volume, dry tissue mass, and body water content

We adapted the protocol previously described in ^25^. Synchronized third instar larvae and newly emerged adults were weighted to obtain their wet mass. Larvae were earlier rinsed in PBS and dried in a Kimwipe paper. Hemolymph was blotted out by piercing the body of the insect lying on a PBS-moistened Kimwipe with a tungsten needle. Each animal was reweighted and the difference between the wet mass and this second weighing was taken as an estimate of hemolymph volume. Bodies were then dried at 60ºC overnight and weighted a third time (dry tissue mass). The body water content was computed by subtracting the dry tissue mass from the wet mass. Values were all normalized to *Lpp>+* control.

### Growth curves

Adult females were allowed to lay eggs for 4h in small plates made of PBS, 2% agar, and 2% glucose. After 26h from the beginning of the oviposition, forty first instar larvae were transferred into a tube with standard fly food and incubated at 25ºC. In order to generate pupariation curves, the time to develop was monitored by counting the number of larvae that underwent pupariation every 2-3h. Subsequently, in five groups of forty synchronized white prepupae for each genotype the number of new emerging adults was scored to create eclosion curves.

### Pupal case volume measurements

2-day-old pupae from vials at a density of forty animals were aligned in a Petri dish and captured under bright light by using the Leica M205 FA microscope. Pupal length (L) and width (W) were obtained with Fiji by measuring the medial line between anterior-posterior, and by measuring along the axial line, respectively. Pupal case volume was calculated in Microsoft Excel by using the formula for prolate spheroid: (π/6)L*W^2^. Values were normalized to *Lpp>+* control to give the “Δ pupal case volume (%).”

### Food intake

Food intake was calculated as described in ^26^. Synchronized pre-wandering third instar larvae were washed in PBS, dried on a Kimwipe paper, and incubated for 3h at 25ºC in a petri dish with regular food supplemented with 1.5% w/v of blue dye (Erioglaucine Disodium Salt, Merck, #861146). Larvae were then rinsed in PBS, dried, weighed, immediately frozen in liquid N_2_, and stored at -20ºC. Animals were homogenized in 20μl PBS and centrifuged for 20min at 4ºC. 10μl of supernatant were transferred in a new tube and the amount of blue dye was measured with a spectrophotometer (OD:629nm). Values were next normalized to the larval mass and relative Δ food intake shows variations in food intake in *Lpp>Lsp1α*^*RNAi*^*+Lsp2*^*RNAi*^ and *Lpp>Fbp1*^*RNAi*^ larvae compared to *Lpp>+* control, respectively.

### Wing imaginal disc volume

Wing discs of larvae at 120h AEL were dissected in ice-cold PBS, fixed in 4% formaldehyde for 20min, and mounted in Vectashield Antifade Mounting Medium with DAPI (Vector Laboratories, #H-1200-10) on *Cellview* cell culture dishes with glass bottom without coverslip, in order to preserve the original structural volume. Imaging was performed with a Zeiss LSM900 Inverted Laser Scanning Confocal Microscope and the confocal Z-stacks were processed with the Imaris software (Oxford Instruments) to generate surfaces that represent the original disc volume ^27^. Values were normalized to *Lpp>+* control.

### Motility assay

To test the motility of adult flies, groups of ten animals (five males and five females) were transferred into polystyrene tubes and allowed to acclimatize for 15 to 20 minutes. The tubes were then tapped down on the surface of the bench a few times so that all the flies ended up at the bottom of the tubes. Finally, the time taken for all the animals in the tubes to cross a line placed at 10cm from the bottom was measured. The measurements were repeated at least 10 times per group, and the average climbing speed of the flies of different genotypes was calculated.

### Muscle staining

Tissues were dissected in PBS and immediately fixed 20min in 4% formaldehyde (ThermoFisher Scientific, #28908) at room temperature, washed twice for 5min each in PBS, twice for 5min each in PBST (PBS + 0.3% Triton X-100), blocked for 1h in blocking solution (PBST + 2% BSA), incubated with Alexa Fluor™ 647 Phalloidin (ThermoFisher Scientific, #A22287) overnight at 4ºC and washed five times for 20min each in PBS at room temperature. Adult muscles were mounted in Vectashield Antifade Mounting Medium with DAPI. Finally, images were taken under an Inverted Laser Scanning Confocal Microscope with Spectral Detection (LSM900-Zeiss) and processed with Fiji software.

### Fertility measurements

The egg-laying capacity of *Lpp>+, Lpp>Lsp1*α^*RNAi*^*+Lsp2*^*RNAi*^ and *Lpp>Fbp1*^*RNAi*^ females was assessed by measuring the average number of eggs laid in 24h by a single female crossed with *w*^*1118*^ males. In contrast, male fertility was estimated by crossing them with *w*^*1118*^ virgins and counting the number of eggs hatched. In both cases, ten groups of twenty females and ten males were analysed for each individual cross.

### Adult survival experiments

In order to generate age-synchronized adult flies, larvae were raised on standard food at low density (forty larvae per vial), emerged adults were transferred to new vials and immediately mated. Regarding fly resistance to starvation, adults are transferred immediately after eclosion into tubes containing only agar and PBS. Conversely, lifespan was assessed by rearing flies of the three different genotypes on standard food, transferring to fresh food vials three times per week, and scoring deaths every day. For each genotype at least five vials containing twenty females and ten males were analysed.

### Adult wing area

Adult female flies of the appropriate genotype were collected and stored in ethanol. Wings were dissected and mounted in a lactic acid: ethanol (6:5) solution. Pictures were acquired with a 1024×768 resolution using a MZ16-FA Leica Fluorescence Stereomicroscope with a DFC-490 Leica digital camera (bright-field mode, 50% illumination intensity, 10.5 exposure, 2.3 gain, 152 saturation and 1.2 gamma). Wing area was measured by using an automated deep learning-based segmentation technique previously described ^27^. Values were normalized to *Lpp>+* control.

### Scaling relationships

For the purpose of analyse the wing-body scaling relationship, flies were crossed, and females allowed to oviposit in plates containing PBS, agar, and glucose for 4h. After 26h from the beginning of the oviposition, groups of forty L1 larvae were transferred in new vials with fresh standard food and incubated at 25ºC. Once adult flies of the accurate genotype emerged from pupal case, males and females were immediately separated and incubated at 25ºC for few hours. Flies were then individually weight as described in the paragraph “Wet mass, hemolymph volume, dry tissue mass, and body water content” and wings were dissected and mounted as explained previously. Data were plotted on a log-log scale with wing area on y-axis and body mass on x-axis to make the relationship linear. As previously reported, this relationship can be described using the simple linear equation: log(y) = α log(x) + log(b), where x is body size, y is organ size, log(b) is the intercept of the line on the y-axis and α is the slope of the line, also known as the allometric coefficient ^28^. Equations and significance between the slopes were then calculated with GraphPad Prism.

### Statistics

Statistical analyses were performed with GraphPad Prism V9 and Microsoft Excel V16.74. For each experimental condition, control samples were carried out in parallel with the same set-up. p values and significance were as follows: ns=p>0.05; *=p≤0.05; **=p≤0.01; ***=p≤0.001; and ****=p≤0.0001. For RT-PCRs in Fig. 1A-D and Fig. S1A-F, individual replicates were shown (with means connected only in Fig. 1A-D); n=3. In Fig. 1F-I individual values and means with SEM were presented in scatter plot, while in Fig. S1G-M individual values and means with SEM were presented in interleaved scatter with bars; n=3, significance was determined by unpaired nonparametric t test (Mann-Whitney) with the corresponding control. Ecdysone levels in Fig. S1N were shown in bar graph with means and SEM; n=3, significance was analysed by unpaired t test (Mann-Whitney).

Quantification of western blots was represented by individual replicates and trends over multiple experiments in Fig. 2D-I; n=3, significance was determined by unpaired t test (Mann-Whitney). In Fig.3I individual replicates were reported in bar graph; n=3, significance was evaluated by one-way ANOVA followed by Dunnem’s multiple-comparisons test. Body measurements in Fig. 3A-C, Fig. S3B, D, and Fig. S4A-C were shown as percentage difference from *Lpp>+* controls; individual replicates with means and SEM were analysed by one-way ANOVA followed by Dunnem’s multiple-comparisons test or unpaired t test (Mann-Whitney); n ranges between 20 and 58. For the developmental assays (pupariation and eclosion curves) in Fig. 3D, Fig. S3C, and Fig. 4A means with SEM at different time points are connected by lines; n=3 means average of three vials per genotype in one experiment; one-way ANOVA followed by Dunnem’s multiple-comparisons test was used. Wing disc volume in Fig. 3F were shown as percentage difference from *Lpp>+* control; individual replicates with means and SEM were analysed by one-way ANOVA followed by Dunnem’s multiple-comparisons test; n=21. Data showing climbing speed (Fig. 4B) and fertility (Fig. 4D) were individual replicates with means and SEM analysed by one-way ANOVA followed by Dunnem’s multiple-comparisons test or unpaired t test (Mann-Whitney); n=10 in Fig. 4B means the average of ten different measurements of climbing speed for ten groups of flies; n=10-19 in Fig. 4D means the average of eggs laid or hatched in 10-19 group of flies. Adult wing area measurements in Fig. S4D were shown as percentage difference compared to corresponding controls; n=24-59, significance was tested by one-way ANOVA followed by Dunnem’s multiple-comparisons test. Results regarding survival during starvation (Fig. 4E) are shown in percent staircase graph; n=10 animals, significance was determined by one-way ANOVA followed by Dunnem’s multiple-comparisons test. For lifespan experiments shown in Fig. 4F, n ranges between 60 and 155 females and between 30 and 60 males; results are present as percent staircases with 95% confidence interval, significance was estimated by Log-rank (Mantel-Cox) and Gehan-Breslow-Wilcoxon tests. For static allometry in Fig. 4G and Fig. S4E, measurements were transformed in logarithm and plotted in log-log graph. Equations of simple linear regression and tests to check if slopes and intercepts differ significantly were generated by GraphPad Prism. n=20 means twenty females and twenty males per genotype were analysed.

## ACKNOWLEDGMENTS

We thank Stephen Simpson, Allison Bardin, Yohanns Bellaïche and members of the laboratory for discussions and comments on the manuscript; S. Eaton laboratory for kindly providing *Lpp-Gal4* strain and Cv-d antibody; N. Tapon for *UAS-TSC1+TSC2* flies; the Bloomington Stock Center and the Vienna *Drosophila* Resource Center for fly stocks; BestGene for generating the *QUAS-Lsp1α-HA* and *QUAS-Lsp2-HA* fly lines; Eurogentec for antibody production; and the PICT-IBiSA@BDD light-microscopy facility of Institut Curie. This work was supported by Institut Curie, CNRS, INSERM, FRM, European Research Council (Advanced Grant n°694677 to P.L.), Labex DEEP program (ANR-11-LABX-0044, ANR-10-IDEX-0001-02).

## AUTHOR CONTRIBUTIONS

Conceptualization, L.V. and P.L.; Methodology, L.V., A.A. and P.L.; Investigation, L.V.; Writing – Original Draj, L.V.; Writing – Review & Editing, L.V. and P.L.; Funding Acquisition, P.L.; Supervision, L.V. and P.L.

