## Supplementary material for "A temporal allocation of amino acid resources ensures fitness and body allometry": Suppl Figures

### SUPPLEMENTARY FIGURES

#### Figure S1

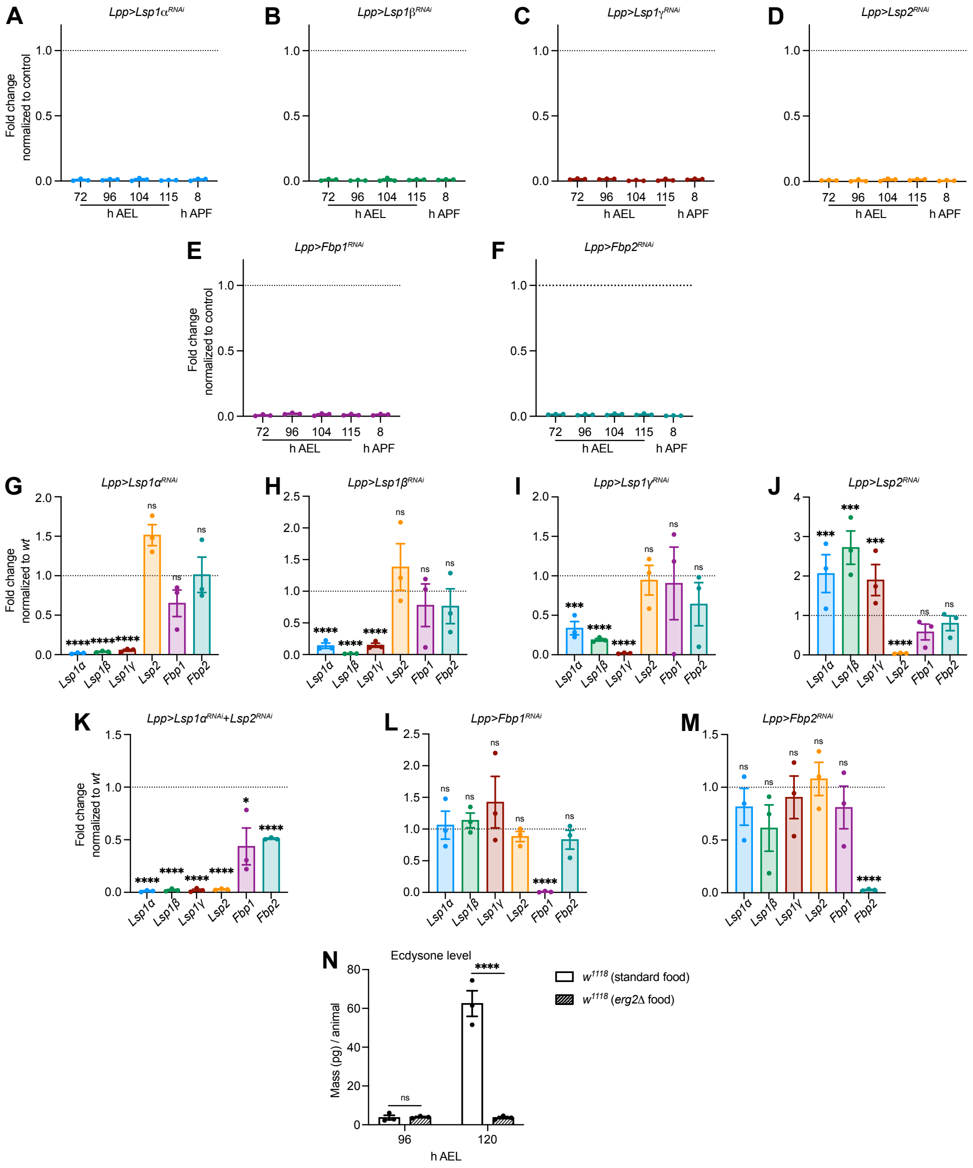


**Figure S1. The fat body is the major site of LSP and FBP production. Related to Figure 1.** (A-F) Total body mRNA levels of *Lsp1α* (A), *Lsp1β* (B), *Lsp1γ* (C), *Lsp2* (D), *Fbp1* (E), and *Fbp2* (F) during the development, measured by RT-PCR, in larvae in which one of the *Lsp* or *Fbp* gene was downregulated only in the fat body. Results shown are mean±SEM and n=3 replicates per time point. Fold changes are normalized by *rp49* to *Lpp>+* control larvae; developmental time points are expressed in hours after egg laying [h AEL], and hours after pupa formation [h APF].

(G-M) Total body mRNA levels of *Lsp1*$\alpha$, *Lsp1*$\beta$, *Lsp1*$\gamma$, *Lsp2*, *Fbp1*, and *Fbp2* in larvae in which one gene of the LSPs-FBPs system was silenced, measured by RT-PCR. Results shown are mean±SEM and n=3 replicates for each gene analyzed. Fold changes are normalized by *rp49* to *Lpp>+* control larvae (unpaired nonparametric t test (Mann-Whitney). ns: not significant, *p≤0.05, ***p≤0.001, ****p≤0.0001).

(N) Quantification of the circulating 20-Ecdysone in larvae reared on standard or in *erg2*$\Delta$ food. Larvae were bled at 96h and 120h after egg laying (n=3 replicates per time point. Unpaired nonparametric t test (Mann-Whitney). ns: not significant, ****p≤0.0001).

### Figure S2

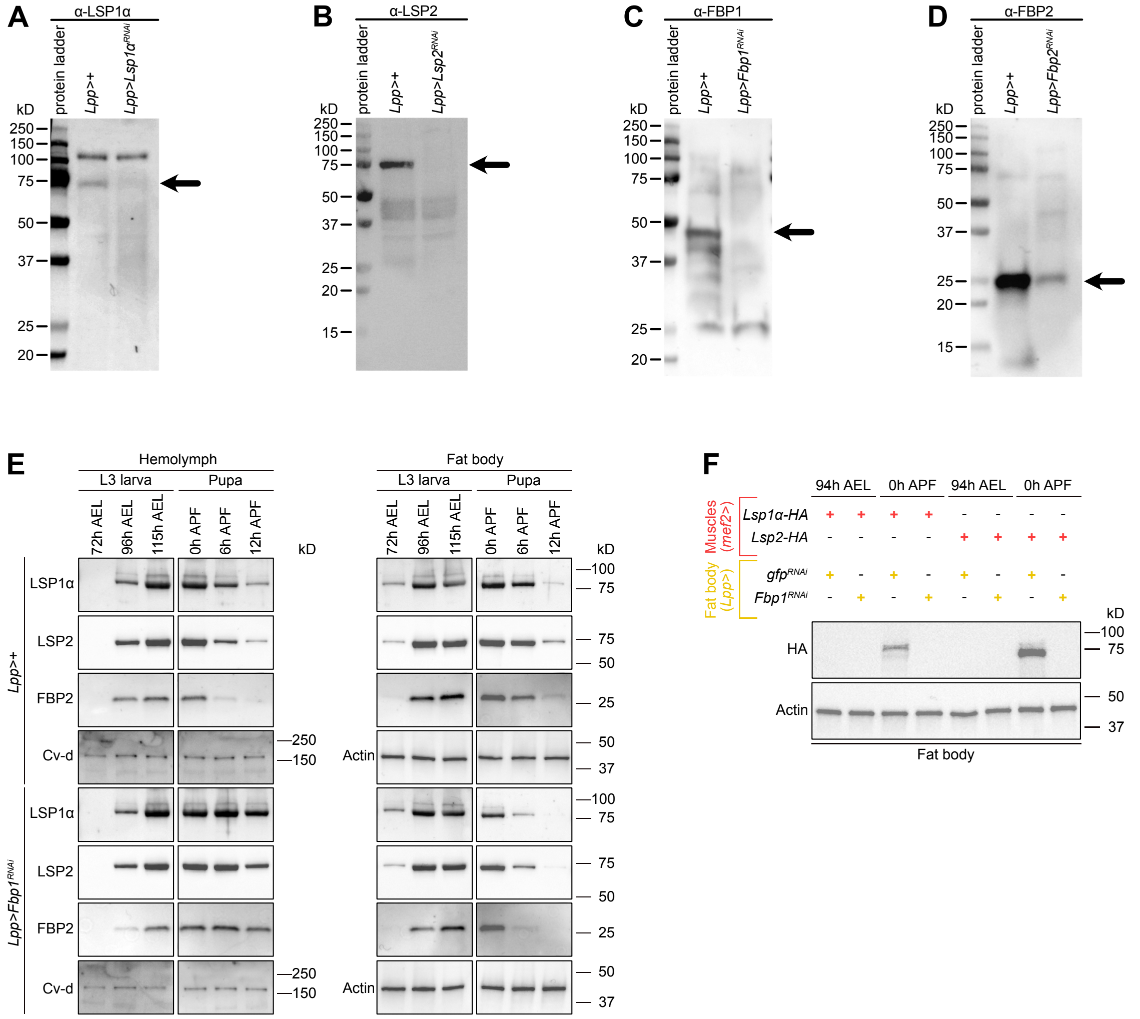


**Figure S2. FBP1 promotes the internalization of LSPs in the fat body. Related to Figure 2.**

(A-D) Immunoblots of whole-body protein extracts from late wandering larvae of the indicated genotypes. Samples were probes with antibodies to LSP1α (A), LSP2 (B), FBP1 (C), and FBP2 (D) to test their efficacy. Molecular masses are reported on the left and a black arrow indicates the band corresponding to the target protein.

(E) Immunoblots of hemolymph and fat body extracts from larvae and pupae with the indicated genotypes. Samples were collected at different hours after egg laying [h AEL] and hours after pupa formation [h APF], and they were probed with antibodies to LSP1α, LSP2, and FBP2. Cv-d and Actin antibodies were used as loading control for hemolymph and fat body respectively. Molecular masses are indicated to the right of each blot. Quantification of the bands is shown in Figure 2D-I.

(F) Immunoblots of fat body extracts from larvae and pupae at 94h AEL and 0h APF expressing different transgenes in muscles (indicated in red) and in fat body cells (reported in yellow). Samples were probed with antibodies against HA and Actin, with molecular masses indicated to the right.

### Figure S3

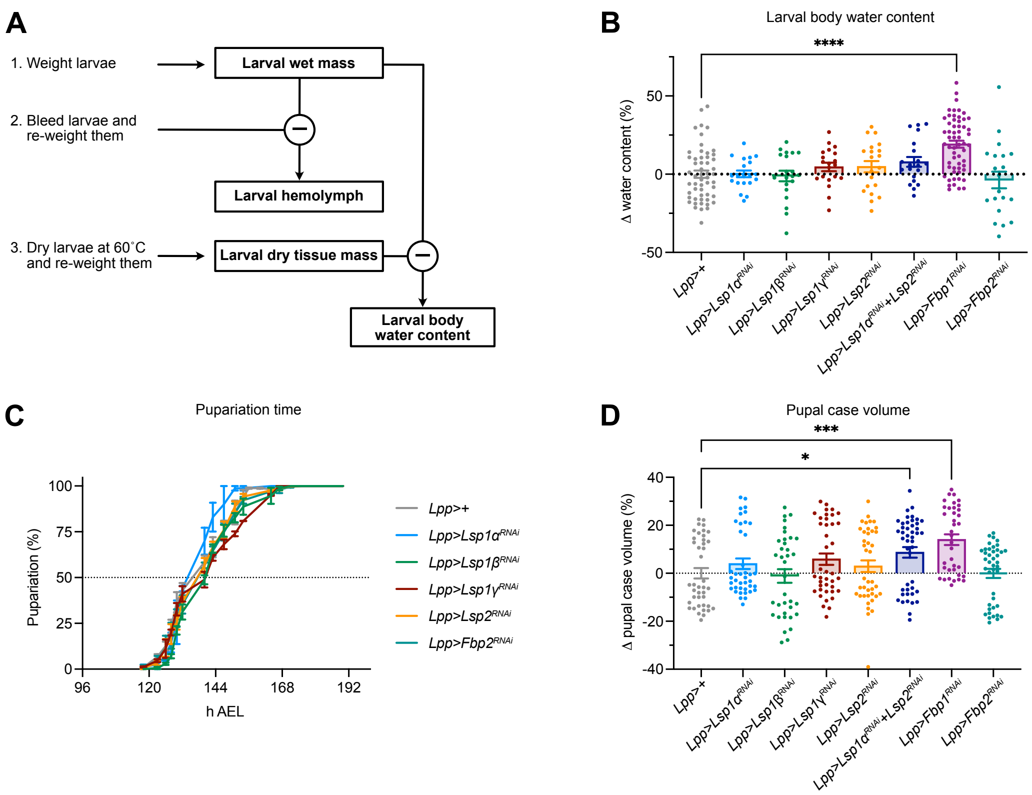


**Figure S3. Hexamerin absence impairs larval body growth. Related to Figure 3.**

(A) Flowchart of the protocol used in this study to analyze various body parameters.

(B) Quantification of larval water body content with the indicated genotypes (n=20-58 for each genotype). Results shown are mean±SEM as percentage difference compared to *Lpp>+*. Larvae of all genotypes were compared to control, but only the significantly different pairs are reported in the graph (one-way ANOVA followed by Dunnett's multiple-comparisons test. ****p≤0.0001).

(C) Pupariation timing of larvae with the indicated genotypes. Means with SEM at different time points are connected by lines; n=3 means average of three vials per genotype in one experiment. All curves were compared to *Lpp>+* one using one-way ANOVA followed by Dunnett's multiple-comparisons test: none of them shown significant difference.

(D) Variations in pupal case volume are represented in the graph (n=36-45 for each genotype). Percentage values are the difference between *Lpp>+* control pupae and the analyzed genotypes (one-way ANOVA followed by Dunnett's multiple-comparisons test. Pairs showing significant difference are indicated in the graph. *p≤0.05, ***p≤0.001).

### Figure S4

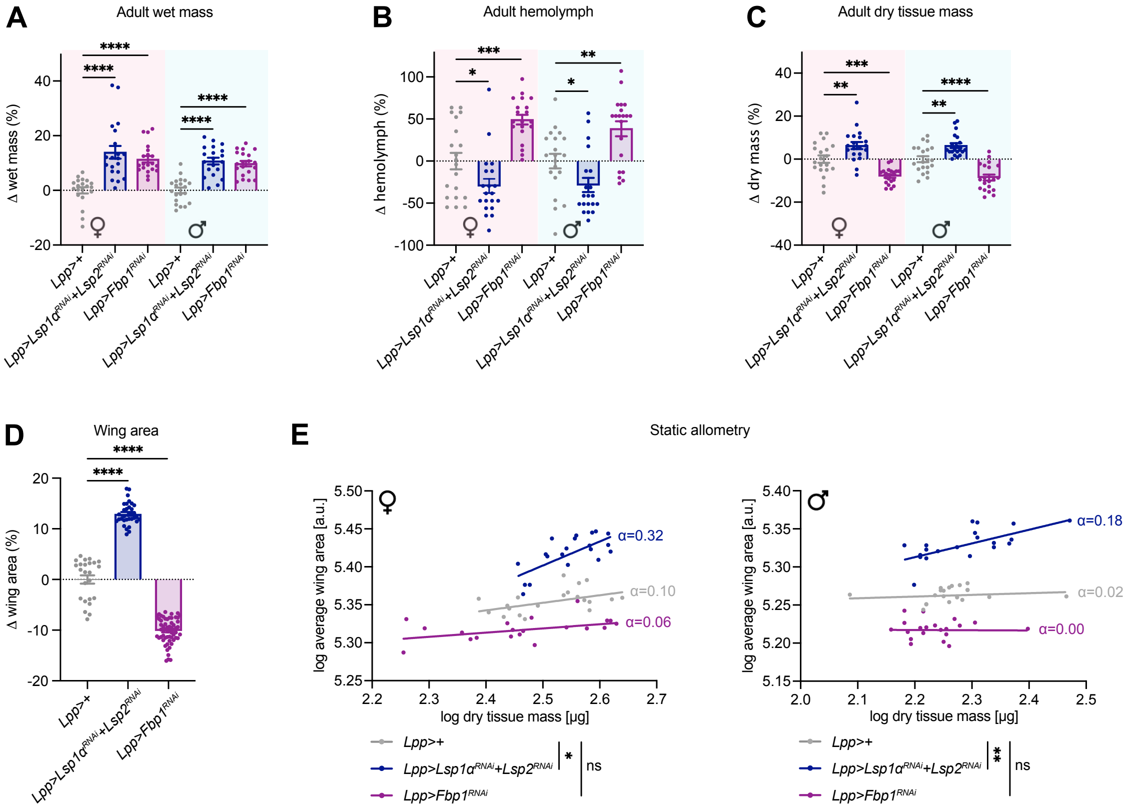


**Figure S4. Consequences of amino acid storage dysfunction on adult body. Related to Figure 4.**

(A-C) Quantification of different body parameters with the indicated genotypes (n=20 females and 20 males for each genotype and time point). Females and males were measured at 0h AE [after eclosion]. In all graphs results shown are mean±SEM as percentage difference compared to control *Lpp>+* (one-way ANOVA followed by Dunnett's multiple-comparisons test. *p≤0.05, **p≤0.01, ***p≤0.001, ****p≤0.0001).

(D) Variation of the wing area between the control *Lpp>+*, *Lpp>Lsp1*$\alpha^{RNAi}$*+Lsp2*$^{RNAi}$ and *Lpp>Fbp1*$^{RNAi}$ females (n=24-59 per genotype). Results are shown as percentage difference compared to control *Lpp>+* (one-way ANOVA followed by Dunnett's multiple-comparisons test. ****p≤0.0001).

(E) The scaling relationship for wing size against dry tissue mass was obtained by transforming raw data in logarithm and plotted in log-log graph; n=20 females and 20 males per genotype (equations for simple linear regression and tests for slope and intercept significancy were generated by GraphPad Prism. *p≤0.05, **p≤0.01). Allometric coefficients ($\alpha$) of the different lines are reported in the graphs.
